# Network structure of the human musculoskeletal system shapes neural interactions on multiple timescales

**DOI:** 10.1101/181818

**Authors:** Jennifer N. Kerkman, Andreas Daffertshofer, Leonardo L. Gollo, Michael Breakspear, Tjeerd W. Boonstra

## Abstract

Human motor control requires the coordination of muscle activity under the anatomical constraints imposed by the musculoskeletal system. Interactions within the central nervous system are fundamental to motor coordination, but the principles governing functional integration remain poorly understood. We used network analysis to investigate the relationship between anatomical and functional connectivity amongst 36 muscles. Anatomical networks were defined by the physical connections between muscles and functional networks were based on intermuscular coherence assessed during postural tasks. We found a modular structure of functional networks that was strongly shaped by the anatomical constraints of the musculoskeletal system. Changes in postural tasks were associated with a frequency-dependent reconfiguration of the coupling between functional modules. These findings reveal distinct patterns of functional interactions between muscles involved in flexibly organising muscle activity during postural control. Our network approach to the motor system offers a unique window into the neural circuitry driving the musculoskeletal system.

## Introduction

The human body is a complex system consisting of many subsystems and regulatory pathways. The musculoskeletal system gives the body structure and creates the ability to move. It is made up of more than 200 skeletal bones, connective tissue and over 300 skeletal muscles. Muscles are attached to bones through tendinous tissue and can generate movement around a joint when they contract. The central nervous system controls these movements through the spinal motor neurons, which serve as the final common pathway to the muscles (*1*). While the anatomical and physiological components of the musculoskeletal system are well characterized (*2, 3*), the organisational principles of neural control remain poorly understood. Here we elucidate the interplay between the anatomical structure of the musculoskeletal system and the functional organisation of distributed neural circuitry from which motor behaviours emerge.

The traditional idea that the cortex controls muscles in a one-to-one fashion has been challenged by several lines of evidence (*4*). For example, it is widely recognized that the many degrees-of-freedom (DOFs) of the musculoskeletal system prohibit a simple one-to-one correspondence between a motor task and a particular motor solution; rather muscles are coupled and controlled in conjunction (*5*). A coupling between muscles - whether mechanical or neural - reduces the number of effective DOFs and hence the number of potential movement patterns. This coupling thereby reduces the complexity of motor control (*6*).

There is continuing debate about the nature of the coupling between muscles. The mechanical coupling in the musculoskeletal system constrains the movement patterns that can be generated (*7, 8*). For example, the biomechanics of the limb constrain relative changes in musculotendon length to a low dimensional subspace, resulting in correlated afferent inputs to spinal motor neurons (*9*). The coupling between muscles could also result from redundancies in the neural circuitry that drives spinal motor neurons (*10*). Electrophysiological studies reveal that a combination of only a few coherent muscle activation patterns - or muscle synergies - can generate a wide variety of natural movements (*11*). Some of these patterns are already present from birth and do not change during development, whereas other patterns are learned (*12*). This supports the notion that the neuromuscular system has a modular organisation that simplifies the control problem (*13*). Spinal circuitry consisting of a network of premotor interneurons and motor neurons that may generate basic movement patterns by mediating the synergistic drive to multiple muscles (*14*). These spinal networks may encode coordinated motor output programs (*15*), which can be used to translate descending commands for multi-joint movements into the appropriate coordinated muscle synergies that underpin those movements (*3*).

Network theory can provide an alternative perspective on the modular organisation of the musculoskeletal system. One of the most relevant features of complex networks are community or modular structures, which refer to densely connected groups of nodes with only sparse connections between these groups (*16*). The investigation of community structures has been widely used in different domains such as brain networks (*17*). It has recently been applied to investigate the structure and function of the musculoskeletal system: The anatomical network can be constructed by mapping the origin and insertion of muscles (*18, 19*). We have previously shown how functional muscle networks can be constructed by assessing intermuscular coherence from surface electromyography (EMG) recorded from different muscles (*20*). These functional networks reveal functional connectivity between groups of muscles at multiple frequency bands. Coherence between EMGs indicates correlated or common inputs to spinal motor neurons that are generated by shared structural connections or synchronisation within the motor system (*10, 21, 22*). Functional connectivity patterns hence allow to assess structural pathways in the motor system using non-invasive recordings (*23*).

Here we investigate the organisational principles governing human motor control by comparing the community structure of anatomical and functional networks. We use multiplex modularity analysis (*24*) to assess the community structure of functional muscle networks across frequencies and postural tasks. As biomechanical properties of the musculoskeletal system constrain the movement patterns that can be generated, we expect a similar community structure for anatomical and functional muscles networks. Deviations in community structure indicate additional constraints imposed by the central nervous system. We also compare functional connectivity between modules during different tasks to investigate changes in functional organisation during behaviour. While average functional connectivity is constrained by anatomical constraints, we expect that functional muscle networks reconfigure to enable task-dependent coordination patterns between muscles. Such task modulations would indicate that functional interactions between muscles are not hard-wired but are instead governed by dynamic connectivity in the central nervous system that is shaped by the anatomical topology of the musculoskeletal system.

## Results

We assessed the relationship between anatomical and functional connectivity of key muscles involved in postural control tasks (36 muscles distributed throughout the body). We investigated a muscle-centric network in which the nodes represent the muscles and the edges of the network are anatomical connections or functional relations between muscles.

### Anatomical muscle network

Anatomical muscle networks were defined by mapping the physical connections between muscles (*19, 25*), based on gross human anatomy (*2*). The anatomical network constituted a densely connected, symmetrical network (Fig. 1; network density is 0.27). Modularity analysis revealed five modules that divided the anatomical muscle network into the main body parts (right arm, left arm, torso, right leg and left leg) with a modularity of 0.39.

**Figure 1.**
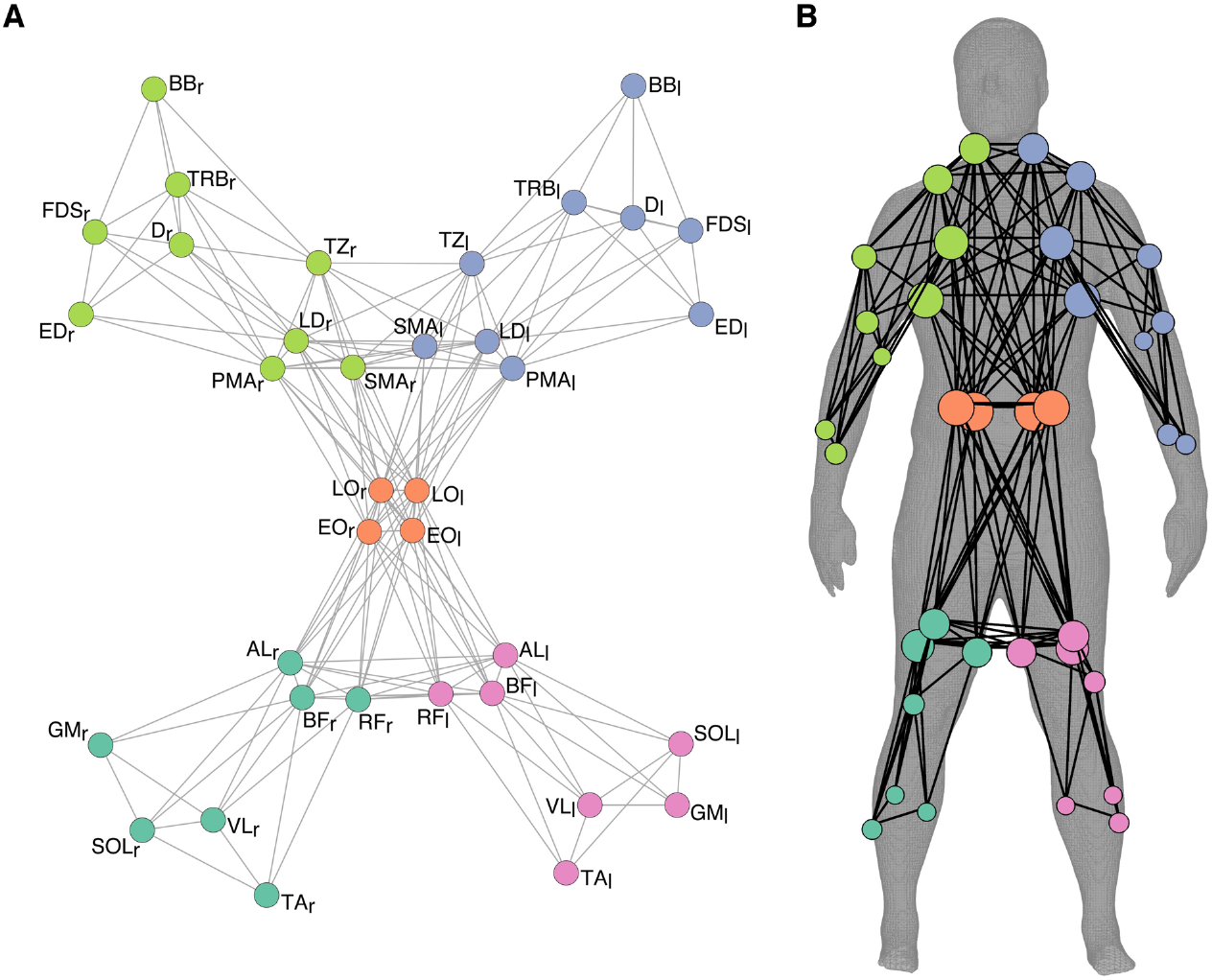
Community structure of the anatomical muscle network. **A)** Topological representation of the anatomical network. The nodes of the network represent the muscles and edges represent anatomical connections between muscles that are attached to the same bone or connective tissue. The five modules are colour-coded. **B)** Spatial representation of anatomical muscle network displayed on the human body (*26*). The size of each node represents the number of other nodes it is connected to.

### Functional muscle network

Functional muscle networks were defined by mapping correlated inputs to different muscles. To map functional networks, we measured surface EMG from the same 36 muscles while healthy participants performed different postural tasks. A full factorial design was used in which we varied postural control (normal standing, instability in anterior-posterior or medial-lateral direction) and pointing behaviour (no pointing, pointing with the dominant hand or with both hands; see Methods for details). We used these tasks to experimentally manipulate the required coordination between muscles and to induce changes in the functional muscle network. We assessed functional connectivity by means of intermuscular coherence between all muscle combinations and used non-negative matrix factorisation (NNMF) to decompose these coherence spectra into frequency components and corresponding edge weights. This yielded a set of weighted networks with their corresponding spectral fingerprints (frequency components).

We observed four separate frequency components (component 1: 0-3 Hz, component 2: 3-11 Hz, component 3: 11-21 Hz, component 4: 21-60 Hz; Fig. 2A), which serve as separate layers of a multiplex network and explained most of the variance of the coherence spectra (*R*^2^ = 0.90). Weights were thresholded to obtain a minimally connected binary network across layers and to keep the number of edges constant across layers (relative threshold of 0.035). Using multiplex modularity analysis, we obtained a fixed community structure across all four frequencies and nine conditions, which revealed six modules: right upper arm (rUA), bilateral forearms (FA), torso (T), right upper leg (rUL), left upper leg (lUL) and bilateral lower legs (LL) (Fig. 2B). Figure 2C depicts how these modules are distributed across the body. Distinct network topologies were observed across layers with a more widely connected network at lower frequencies and more partitioned network at higher frequencies: network density was 0.10, 0.09, 0.08, and 0.06 and the modularity was 0.46, 0.60, 0.64, and 0.75 for components 1 to 4, respectively (Fig. 2D).

**Figure 2.**
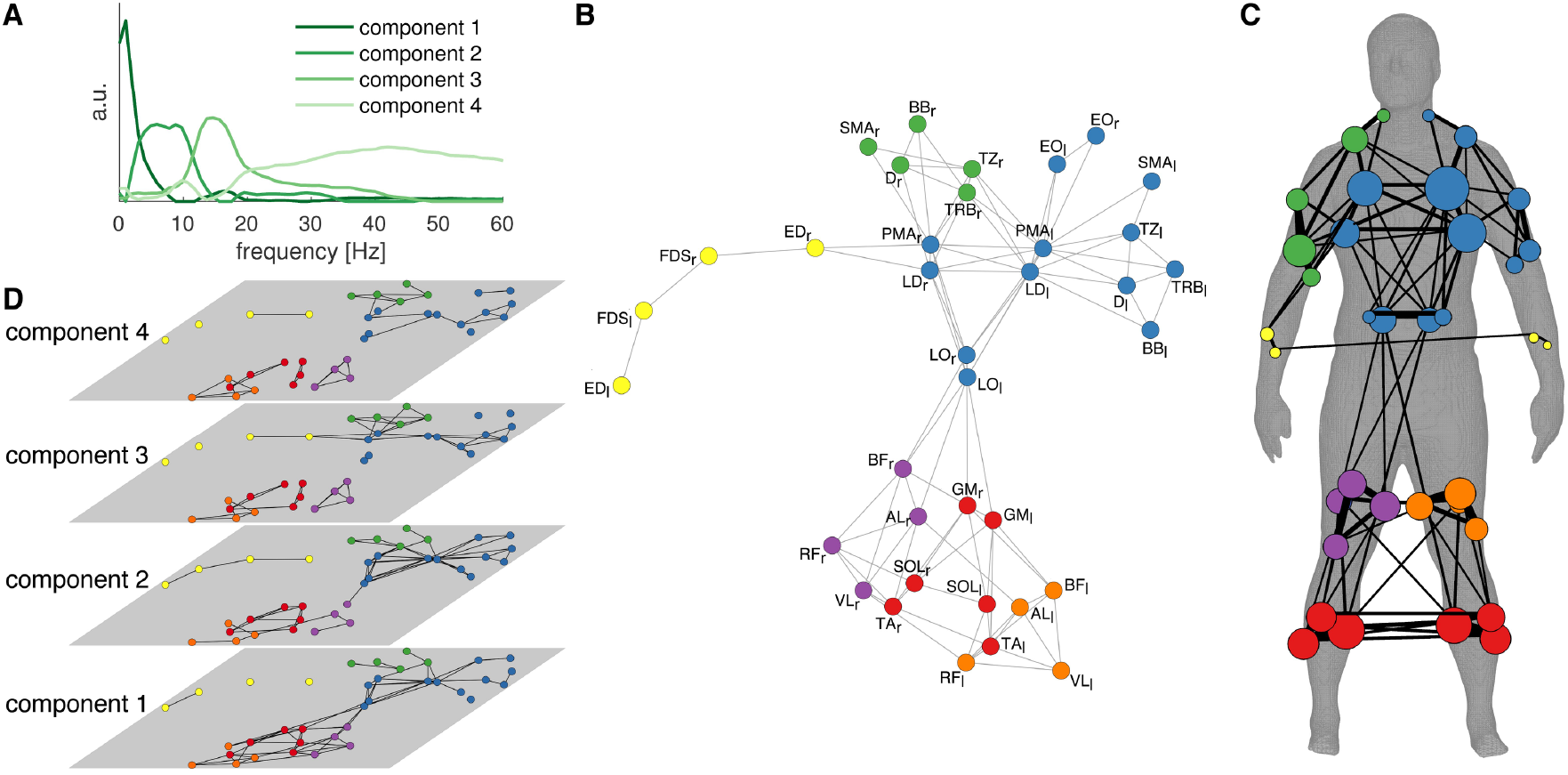
Community structure of multiplex functional muscle networks. **A)** The frequency spectra of the four components obtained using NNMF. **B)** Multiplex community structure of functional muscle network across frequencies and conditions. The dominant hand of all participants is displayed on the right side of the human body. **C)** Spatial representation of the average muscle network displayed on the human body (*26*). The size of the nodes represents the number of other nodes it is connected to and the width of the number of edges across layers. **D)** The binary muscle networks for each layer.

### Comparison between anatomical and functional networks

The community structures of the anatomical and functional muscle networks were very similar (Rand index = 0.80, adjusted Rand index = 0.36, *P* < 0.001). A marked difference between anatomical and functional networks are the connections between bilateral forearm and bilateral lower leg muscles in the functional networks, which were absent in the anatomical network. This is reflected in the community structure of the functional networks, where bilateral lower leg muscles and bilateral forearm muscles were grouped in separate modules (Fig. 2C).

**Figure 3.**
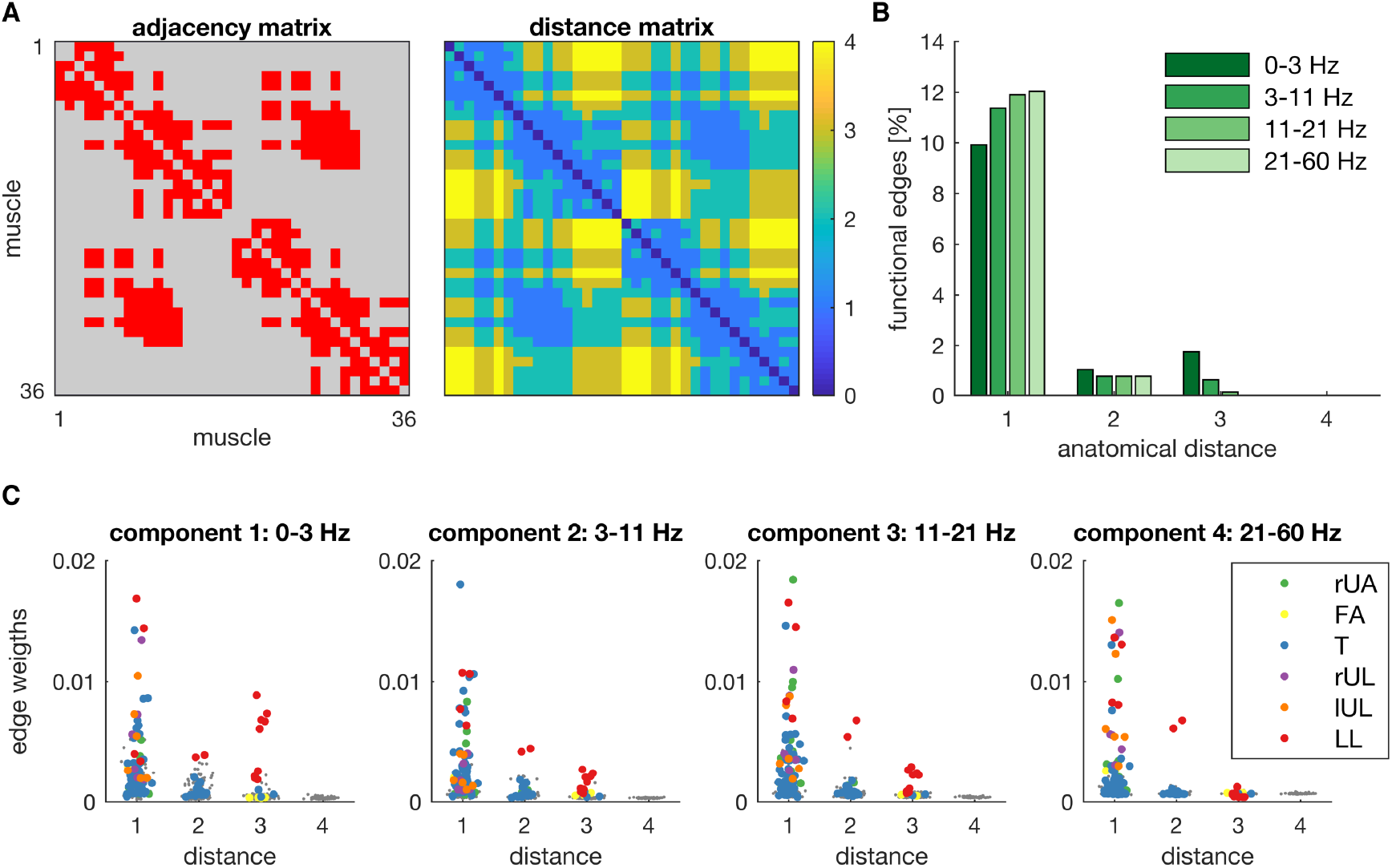
Relationship between functional connectivity and anatomical distance. **A)** Adjacency and distance matrix of the anatomical muscle network. Maximum anatomical distance (path length) is 4. **B)** Percentage of functional edges of thresholded networks across experimental conditions as a function of anatomical distance. **C)** Distribution of edge weights of functional networks as a function of anatomical distance for each layer. Weights were averaged across experimental conditions. Edges connecting muscles within the same module are colour-coded (rUA: right upper arm, FA: bilateral forearms, T: torso, rUL: right upper leg, lUL: left upper leg, and LL: bilateral lower legs) and grey dots represent edges between modules.

The comparison between anatomical distance (path length) and functional connectivity revealed that anatomically nearby nodes are more likely to receive common input (Fig. 3). We first examined the percentage of all possible edges, i.e. the number of edges above threshold, which decreased as a function of anatomical distance: 11.3%, 0.9%, 0.6% and 0.0% for anatomical distance 1 to 4, respectively. This decline with distance was even more pronounced for the higher frequency components (Fig. 3B). Next, we examined the distribution of functional weights as a function of anatomical distance. The highest weights were observed for edges connecting muscles within the same module. The edges within most modules had an anatomical distance of 1. Only a few edges had an anatomical distance of 2 or 3 and all of these edges were contained in the FA and LL modules. In particular, edges connecting bilateral lower leg muscles (LL) showed relative large weights at an anatomical distance of 3 (Fig. 3C).

### Task-dependent modulations

We next sought to study the influence of task on this structure-function relationship. This was achieved by employing clustered graphs to compare functional muscle networks across task conditions. The functional modules identified using the preceding multiplex modularity analysis form the nodes of these clustered graphs. Figure 4A shows the clustered graphs in the nine experimental conditions and for the four frequency components. The clustered graphs were very sparse, as modules have dense within-module connections but sparse connections between nodes in other modules. Most edges were observed between leg muscles modules (LL, rUL and lUL) at the lowest frequency components (0-3 and 3-11 Hz), consistent with the lower modularity scores, in particular when postural stability was challenged by instability in anterior-posterior or medial-lateral direction. Edges between the arm muscle modules (rUA and **FA)** and the torso (T) were mainly observed at the higher frequency components (11-21 and 21-60 Hz) during pointing (unimanual and bimanual).

**Figure 4.**
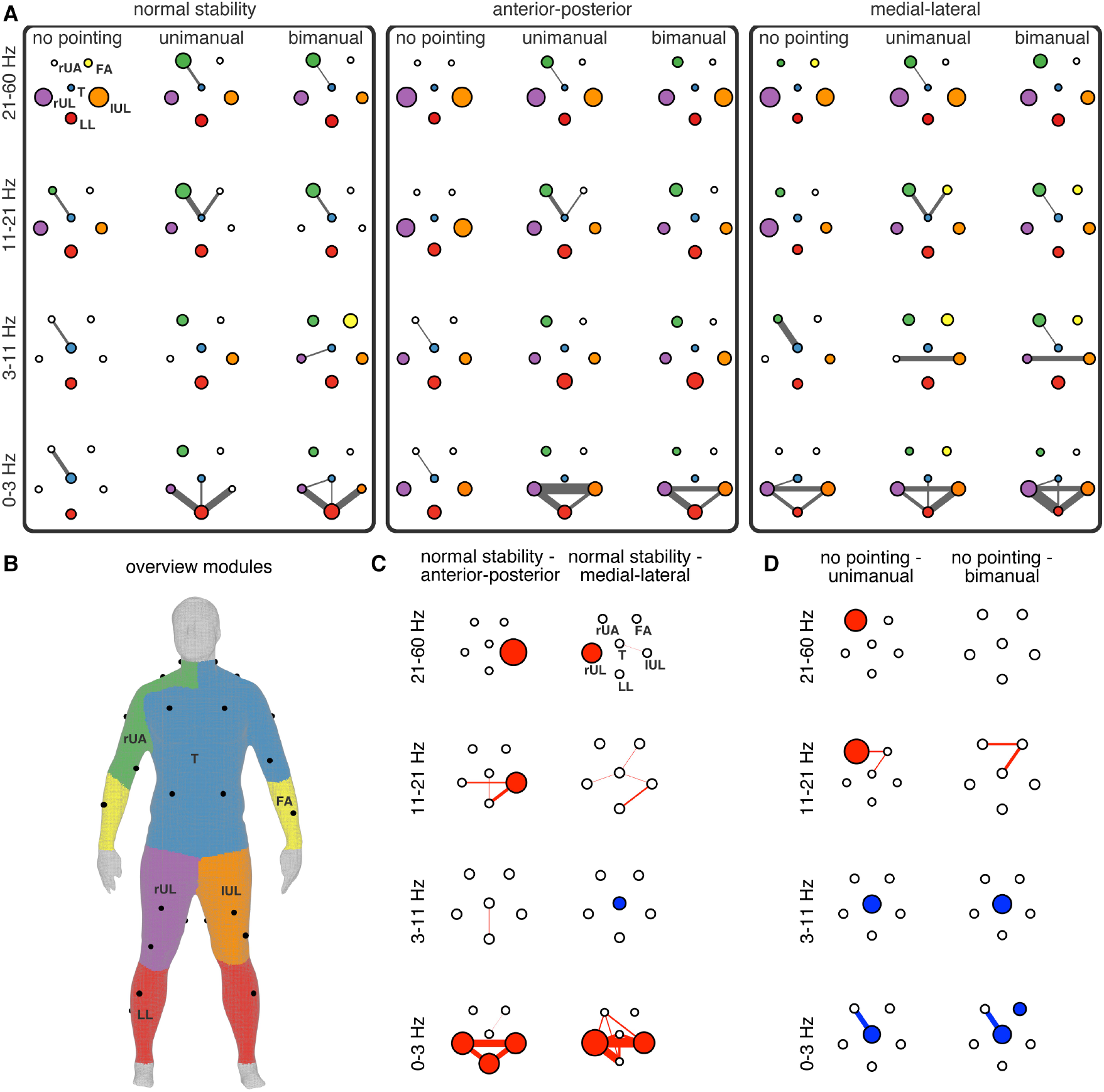
Clustered graphs of functional muscle networks across conditions. **A)** The clustered graphs in the nine experimental conditions (columns) and the four frequency components (rows). The nodes are the modules identified using multiplex modularity analysis. Node size represents the network density within and the width of the edges the connection density between modules. **B)** Spatial representation of the functional modules on the human body: right upper arm (rUA), bilateral forearms (FA), torso (T), right upper leg (rUL), left upper leg (lUL) and bilateral lower legs (LL). We used toolboxes for geometry processing to generate the coloured meshes (*27*) and display it on the human body (*26*). **C)** Significant differences in the connectivity of the clustered graphs between the stability conditions. Two contrasts were assessed: normal stability - anterior-posterior instability and normal stability - medial-lateral instability. A permutation test was used and family-wise error control was maintained using Bonferroni correction (84 comparisons). Significant differences (*P*_corrected_ < 0.05) are colour-coded: Red depicts an increase and blue a decrease in the average weights. Coloured edges and nodes depict significant changes in connectivity between and within modules, respectively. **D)** Significant differences in the connectivity of the clustered graphs between the pointing conditions. Two contrasts were assessed: no pointing - unimanual pointing and no pointing - bimanual pointing.

The effects of the stability tasks were largely confined to the leg modules (Fig. 4C). Increased connectivity was observed during postural instability (anterior-posterior and medial-lateral) compared to normal standing within most frequency components. At the lowest frequency component (0-3 Hz), connectivity increased within and between most leg modules (*P*_corrected_ < 0.01). Only small differences were observed at 3-11 Hz: increased connectivity between the torso (T) and lower leg (LL) modules during anterior-posterior instability (+25%, range [-9, 46%], *P*_corrected_ = 0.01) and decreased connectivity within the torso module during medial-lateral instability (-21%, range [-50, 0.3%], *P*_corrected_ = 0.01). Connectivity increased again at the highest frequency components (11-21 and 21-60 Hz) within and between the torso and leg modules (rUL, lUL, and LL, *P*_corrected_ < 0.02).

The pointing tasks showed a different pattern compared to the postural tasks, but the effects of unimanual and bimanual pointing were very similar (Fig. 4D). During pointing, connectivity decreased within the torso (T) module at the lowest frequency components (0-3 Hz, -61%, range [-90, -1%], *P*_corrected_ < 0.005; 3-11 Hz, - 59%, range [-86, 2%], *P*_corrected_ < 0.02) and between the torso and the right upper arm (rUA) module only at the lowest frequency component (0-3 Hz, -67%, range [-93, -9%], *P*_corrected_ < 0.005). In contrast, a significant increase in connectivity within the rUA module was observed during unimanual pointing compared to no pointing at the highest frequency components (11-21 Hz, +64%, range [-4, 95%], *P*_corrected_ = 0.005; 21-60 Hz,+66%, range [-12, 93%], *P*_corrected_ = 0.015). In addition, there was increased connectivity between the torso and the forearm (FA) modules (+41%, range [-8, 82%], *P*_corrected_ < 0.01) and between rUA and FA (+44%, range [0, 82%], *P*_corrected_ < 0.005) during pointing (unimanual and bimanual) compared to no pointing at frequency component 3 (11-21 Hz).

## Discussion

We used a network approach to study the structure-function relationship of the human musculoskeletal system. Several principles of the functional relationship between muscles were uncovered: (i) Functional connectivity patterns between muscles are strongly shaped by the anatomical constraints of the musculoskeletal system, with functional connectivity strongest within anatomical modules and decreased as a function of anatomical distance; (ii) Bilateral connectivity between the homologous upper and between the homologous lower extremities is a key characteristic of the functional muscle networks; (iii) The functional relationships are task-dependent with postural tasks differentially impacting upon functional connectivity at different frequency ranges. The use of a multiplex approach allows the integration of functional muscle networks across frequencies and provides a unifying window into the distributed circuitry of the human central nervous system that controls movements by innervating the spinal motor neurons.

Identifying relationships between anatomical and functional muscle networks is crucial for understanding how movement is coordinated. Previous studies either investigated how biomechanical properties of the musculoskeletal system constrain the movement patterns that can be generated (*8, 9*), or how muscle activation patterns can be explained by a combination of only a few coherent muscle activation patterns (*11*). Our combined analyses of anatomical and functional muscle networks reveal a strong relationship between the anatomical connections in the musculoskeletal systems and correlated inputs to spinal motor neurons. This builds on previous research showing that common input is strongest to spinal motor neurons that innervate muscles pairs that are anatomically and functionally closely related (*10, 21*). The similarity between structural and functional networks has been a signature of the study of brain networks (*28*) and the topology of brain networks depends on the brain’s spatial embedding (*29*). The present findings suggest that the principles governing embodied structural and functional networks also applies to the neural circuitry that controls movements and may hence reflect a general principle of the central nervous system.

The similarities between anatomical and functional connectivity may indicate that the anatomical structure constrains the functional interactions between muscles. The anatomical connections between muscles remain largely unchanged over the lifespan (*30*) and it is more likely that the fast-changing functional networks are constrained by the much slower changing anatomical networks than vice versa. These constraints may be imposed through afferent activity. The musculoskeletal properties of the human body restrict the postural dynamics (*9*) and these mechanical couplings result in correlated proprioceptive feedback to spinal motor neurons. The influence of biomechanics on functional muscle networks is expected to be most pronounced at the lower frequency components, as muscles act as a low pass filter of neuronal inputs and kinematics of the musculoskeletal system unfold on a slow time scale. This generates correlated activity at low frequencies that are fed back to spinal motor neurons via sensory afferents. The spatial distribution of common input would arguably mirror the topology of the musculoskeletal system.

Anatomical constraints may also be imposed during neural development. During early development, changes in the topographical distribution of axon terminals of descending projects are dependent on patterns of motor activity and anatomical connectivity between muscles (*31*). Likewise, large changes in functional coupling is observed in infants between 9 and 25 weeks, which reflects a sensitive period where functional connections between corticospinal tract fibres and spinal motor neurons undergo activity-dependent reorganization (*32*). The anatomy of the musculoskeletal system will limit the motor activity patterns that can be performed.

Anatomical and functional connectivity between muscles may also both be influenced by external factors. For example, the connectivity patterns of descending pathways is in part genetically determined (*33*). A somatotopic organisation is observed across the neural motor system and the community structure of the anatomical muscle network mirrors the organisation of primary motor cortex control modules (*19*). Likewise, the spatial organisation of motor neurons of the spinal cord is also related to the anatomical organisation of muscles (*34*) and muscles that are anatomically closely located to each other are also innervated by the same spinal nerves (Fig. S2) (*2*). The topographic organisation of spinal motor neurons is similar across species (*35*) and may hence be a result of evolutionary conservation (*36*). Musculoskeletal anatomy and neuronal pathways are hence both subject to some sort of genetic control.

Functional connectivity was not entirely determined by anatomy and we observed several key differences between anatomical and functional muscle networks. Bilateral modules consisting of muscles in the upper or lower extremities were a key characteristic of the functional muscle network that were absent in the anatomical network. The two bilateral forearm muscles (FDS and ED) showed coherent activity at 3-11 Hz, consistent with previous studies showing bimanual coupling at ~10 Hz between homologous hand and forearm muscles (*37, 38*). The observed bimanual coupling at 3-11 Hz may be generated by the olivocerebellar system, which is known to produce oscillations in this frequency range and for its involvement in the formation of functional muscle collectives (*37*). The bilateral forearm muscles were only weakly coupled to other muscles (Fig. 2), which may reflect the relatively high proportion of direct corticospinal projections - and thus a relative low proportion of diverging projections - to motor neurons innervating hand and forearm muscles (*39*).

In contrast, the bilateral module of lower leg muscles revealed strong coupling at multiple frequency bands, consistent with previous analyses on functional muscle networks (*20*), and showed the strongest long-range connections observed in the present study (Fig. 3C). Bilateral connectivity between arm and leg muscles during balancing could be generated by the vestibulospinal tract, which is known to be involved in postural stability and innervate the spinal grey matter bilaterally (*21*). Bilateral connectivity has been observed at all levels of the corticospinal axis (*40*) and is paramount for functional brain networks, particularly between homologous left-right cortical regions (*41*). The present findings suggest that bilateral coupling is also a defining feature of functional muscle networks. The differences in functional connectivity between bilateral arm and bilateral leg muscles indicate that the functional muscle network - like the anatomical muscle network (*25*) - does not show serial homology.

Functional connectivity displayed distinct task-dependent modulations that were linked to the task the subjects performed: functional connectivity was increased within and between the leg modules during postural instability and increased within and between arm and upper body modules in the pointing conditions.

Functional connectivity between muscles is thus task dependent (*21, 38*), which may suggest the presence of multifunctional circuits in which a given anatomical connectivity pattern can generate different functional activity patterns under various conditions (*42*). Such a distributed circuitry creates the substrate to support many behaviours that are driven by the concerted actions of a large distributed network as opposed to simple, dedicated pathways. The underlying network connectivity hence constrains the possible patterns of population activity to a low-dimensional manifold spanned by a few independent patterns - neural modes - that provide the basic building blocks of neural dynamics and motor control (*43*). Again, this finds similarities with recent investigations of the functional principles of cognitive networks in the brain (*44*).

Task-dependent changes occurred at different frequencies, which indicate the functioning of a multiplex network organisation, whereby the four frequency components reflect different types of interactions between muscles. Four distinct frequency components (0-3, 3-11, 11-21, and 21-60 Hz) were extracted using NNMF. These frequency bands closely match those found previously (*20*), demonstrating the robustness of this finding. An interesting possibility is that these frequency components reflect the spectral fingerprints of different pathways that project onto the spinal motor neurons. It has been suggested that these different frequencies may have specific roles in coding motor signals (*45*). Functional connectivity at the lowest frequency components may result from afferent pathways, while functional connectivity at higher frequencies may reflect correlated input from descending pathways. For example, functional connectivity in the beta band (15-30 Hz) most likely reflects corticospinal projections (*10, 38*). The highest frequency components observed in this study (21-60 Hz) showed the most local connectivity patterns. These local connectivity patterns may reflect propriospinal pathways (*3, 15*). These functional connectivity patterns may be used to uncover the contribution of structural pathways in the formation of coordinated activity patterns in the motor system (*23*). These findings mirror observations in cortical networks where frequency-specific networks reveal different topologies and are differentially expressed across brain states (*46*). The differences in the frequency content of functional connectivity observed between the upper limb and lower limb muscles suggests distinct neural circuitry controlling these body parts.

In summary, our network analysis revealed widespread functional connectivity between muscles, indicative of correlated inputs to spinal motor neurons at multiple frequencies. Correlated inputs indicate divergent projections or lateral connections in the neural pathways that innervate spinal motor neurons and can hence be used to assess spinal networks (*23*). These findings are consistent with a many-to-many rather than an one-to-one mapping between brain and muscle (*4*), in which complex movements arise through relatively subtle changes in the coactivation of different distributed functional modes. We present a novel approach that aligns movement neuroscience with current research on brain networks by showing how the central nervous system interacts with the musculoskeletal system of the human body. This approach fits within the broader framework of network physiology, investigating brain-body interactions (*47*). Similar to the current results, research on network physiology has shown that dynamic interactions among organ systems are mediated through specific frequency bands (*48*). We extended this approach by investigating the network topology of functional interactions between muscles, which are mediated through neural pathways within the spinal cord. Future studies may extend the number of muscles that are investigated, include electroencephalography (EEG) to map brain-body networks and investigate the cortical control of muscle networks, and consider individual differences in anatomy.

From a systems biology perspective, the brain and spinal cord are interwoven with the body – they are ‘embodied’ (*7*) – and brain network analysis can thus be extended to investigate the intrinsic organisation of functional networks in the human spinal cord (*49*). Functional interactions between supra-spinal, spinal and peripheral regions can be integrated using network analysis as a common framework. Such an integrated framework is well-placed to provide new insights and interventions for neurological disorders (*50*).

## Material and Methods

### Data acquisition

Fourteen healthy participants (seven males and seven females, mean age 25±8 years, ten right and four left handed) without any neurological, motor disorder or diabetes mellitus and with a BMI below 25 were included in this study. The experiments were approved by the Ethics Committee Human Movement Sciences of the Vrije Universiteit Amsterdam (reference ECB 2014-78) and performed in full compliance with the Declaration of Helsinki. All participants were written and verbally informed about the procedure and signed an informed consent prior to participation.

Participants were instructed to perform nine different postural tasks. A full-factorial design was used in which postural stability (normal standing, instability in anterior-posterior direction and instability in medial-lateral direction) and pointing behaviour (no pointing, pointing with dominant hand, pointing with both hands) were varied. Postural stability was manipulated using a balance board with one degree of freedom, which allowed movement either in the anterior-posterior or medial-lateral direction. In the pointing task, participants held a laser pointer with their dominant hand (unimanual) or with both hands (bimanual) and pointed it on a white target (25 cm^2^) located at a distance of 2.25 meter, parallel to the transversal axis of the body at the height of the acromion of the participant. The experiment hence consisted of nine (3’3) experimental conditions. The duration of a trial was 30 seconds and each condition was repeated six times.

**Table 1.** List of muscles

| **Muscle** | **Abbreviation** |
| --- | --- |
| tibialis anterior | TA |
| gastrocnemius caput mediale | GM |
| soleus | SOL |
| rectus femoris | RF |
| biceps femoris | BF |
| vastus lateralis | VL |
| adductor longus | AL |
| obliquus externus abdominis | EO |
| pectoralis major | PMA |
| sternocleidomastoideus | SMA |
| longissimus | LO |
| latissimus dorsi | LD |
| trapezius | TZ |
| deltoideus | D |
| biceps brachii | BB |
| triceps brachii | TRB |
| extensor digitorum | ED |
| flexor digitorum superficialis | FDS |

Bipolar surface EMG was recorded from 36 muscles distributed across the body (18 bilateral muscles, Table 1). We selected a representative group of antagonistic muscle pairs involved in postural control that can be properly measured with surface EMG due to their location and size. EMG was acquired using three 16-channel Porti systems (TMSi, Enschede, The Netherlands), online high-pass filtered at 5 Hz and sampled at 2 kHz.

**Table 2.** Origin and insertion of muscles

| Muscle | Origin 1 | Origin 2 | Origin 3 | Insertion 1 | Insertion 2 |
| --- | --- | --- | --- | --- | --- |
| TA | tibia |  |  | os cuneiforme mediale | ossa metatarsi |
| GM | femur |  |  | calcaneus |  |
| SOL | fibula | tibia |  | calcaneus |  |
| RF | os coxae<sup>1,*</sup> | os ilium<sup>1,*</sup> |  | tibia |  |
| BF | femur | os ischii<sup>1,*</sup> |  | fibula | tibia |
| VL | femur |  |  | tibia |  |
| AL | os pubis<sup>1,*</sup> |  |  | femur |  |
| EO | costae<sup>2,*</sup> |  |  | linea alba* | os ilium<sup>1,*</sup> |
| PMA | clavicula | costae<sup>2,*</sup> | sternum<sup>2,*</sup> | humerus |  |
| SMA | clavicula | sternum<sup>2,*</sup> |  | os temporale<sup>3,*</sup> |  |
| LO | ligamentum sacrospinale<sup>1,*</sup> | vertebra* |  | costae<sup>2,*</sup> | vertebra* |
| LD | costae<sup>2,*</sup> | fascia thoracolumbalis* | vertebra* | humerus |  |
| TZ | ligamentum nuchae* | os occipitale<sup>3,*</sup> | vertebra* | clavicula | scapula |
| D | clavicula | scapula |  | humerus |  |
| BB | scapula |  |  | radius |  |
| TRB | humerus | scapula |  | ulna |  |
| ED | humerus |  |  | ossa digitorum |  |
| FDS | humerus | radius | ulna | ossa digitorum |  |

<sup>1</sup> Part of the pelvis; <sup>2</sup> Part of the skeleton thoracis; <sup>3</sup> Part of the ossa cranii; \* Connective structure on the midline of the body connecting bilateral muscles. Origin and insertion are based on gross human anatomy as described in Martini, Timmons and McKinley (2), hence ignoring potential individual differences between participants.

### Anatomical muscle network

The anatomical muscle network was defined by mapping the physical connections between muscles. The nodes represent the 36 muscles (18 left and 18 right) and the edges of the network represent the tendinous attachments of muscles onto bones and connective tissue. The structural connections were defined based on the origin and insertion of the muscles (*2*). Bones that show no or almost no motion in the joint between them were considered as one rigid bony structure, i.e. the pelvis, skeleton thoracis or ossa cranii (*25*). The connections between muscles and bones listed in Table 2 denote a bipartite network *C* with muscles as one group and bones as the second group. We then created a muscle-centric network as the one-mode projections of *C*: *B* = *CC^T^* (*19*). This gave a weighted adjacency matrix where the weights reflect the number of attachments by which two muscles are connected. We converted this to a binary network by setting all none-zero weights to 1.

### Functional muscle network

We mirrored the data of the left handers to create a dominant and non-dominant side. EMG data were pre-processed to remove movement and electrocardiography (ECG) artefacts. EMG was band-pass filtered (1-400 Hz) and independent component analysis was used to remove ECG contamination (*51*). One or two independent components were removed for each participant. EMG data was then high-pass filtered (20 Hz) to remove low-frequency movement artefacts. After preprocessing, EMG envelopes were extracted by means of the Hilbert amplitude (*22*).

We followed the procedure described in Boonstra *et al*. (*20*) to extract functional muscle networks from surface EMG. First, complex-valued coherency was estimated and averaged over trials within each condition for each participant. The absolute value of coherency was squared to obtain magnitude-squared coherence. Intermuscular coherence was assessed between all 630 muscle pairs. Next, nonnegative matrix factorisation (NNMF) was used to decompose these coherence spectra across all muscle combinations, conditions and participants into four distinct frequency components and the corresponding weights. This yielded a set of weights for each frequency component, which defined an undirected weighted network for each condition and participant.

These functional networks were converted to binary networks to facilitate comparison to the anatomical network. Weights were thresholded to obtain a minimally connected network across conditions and frequency components. This thresholding procedure yields a single, unique threshold value, which corresponds to the percolation threshold (*52*). This resulted in sparse networks in which each node was connected to at least one other node by an edge at one of the layers of the multiplex network.

### Community structure

The Louvain algorithm was used to extract the modules from the anatomical networks. As the Louvain algorithm is stochastic, we used consensus clustering to obtain a stable partition across 1000 iterations (*53*). Multiplex modularity analysis (*24*) was used to identify the modules of functional muscle network across the conditions and frequency components. We used *MolTi,* a standalone graphical software, to detect communities from multiplex networks by optimising the multiplex-modularity with the adapted Louvain algorithm (https://github.com/gilles-didier/MolTi). Modules were extracted across the 36 (9’4) binary networks. We used the Rand index and the adjusted Rand index to compare the modules of the anatomical and functional muscle networks (*16*).

### Comparison of functional networks across conditions

To facilitate the comparison of functional networks across task conditions, we coarse-grained the networks (*54*). We used the set of functional modules estimated across conditions and frequency components as a frame of reference to coarse-grain the 36 binary networks and then compared the strength of the inter- and intra-module connections across networks using these module boundaries. In the clustered networks the nodes represent the modules (groups of muscles, identified above) and the edges represent the connections between modules. The nondiagonal elements of the resulting weighted adjacency matrix represent the average edge weights between two modules and diagonal elements the average edge weights within a module.

To compare the clustered networks across conditions, we used simple contrasts between task conditions and quantified differences in the numbers of connections between and within modules. We tested four contrasts: (i) unimanual and (ii) bimanual pointing compared to no pointing and (iii) anterior-posterior and (iv) medial-lateral instability compared to normal standing. To test the statistical significance of these contrasts, we performed paired permutation tests separately on each of the matrix elements (*54*). The clustered networks had a much-reduced dimensionality compared to the original functional muscle networks (21 instead of 630 edges). Family-wise error control was maintained using Bonferroni correction to correct for multiple comparisons (4 ’ 21 = 84 comparisons).

## Acknowledgment

We thank Daniele Marinazzo, Simon Farmer, Sjoerd Bruijn, Henk Schutte and Nadia Dominici for their insightful comments. This work was supported by the Netherlands Organization for Scientific Research (NWO 45110-030 and 016.156.346), the ARC Centre of Excellence in Integrative Brain Function, and the Australian National Health and Medical Research Council (APP1037196, APP1110975). The funders had no role in study design, data collection and analysis, decision to publish, or preparation of the manuscript.

## Author contributions

J.K., A.D. and T.B. conceived the study; all authors designed the experiments; J.K. acquired the EMG data; J.K. and T.B. performed the evaluation of the anatomical network; J.K., L.G. and T.B. performed the evaluation of the functional network; T.B. performed the statistical analyses; J.K. and T.B. wrote the manuscript; and all the authors edited the manuscript.

## Data availability

The EMG data used in this study is available at Zenodo: http://doi.org/10.5281/zenodo.1185196.

