## Supplementary Materials for "Network structure of the human musculoskeletal system shapes neural interactions on multiple timescales" (from another version of this preprint: version 3, 2018-03-26)

**Alternative anatomical muscle network**

The anatomical muscle network was defined by mapping the physical connections between muscles. The physical connections were based on origin and insertion described in Martini *et al.* (*2*). We included attachments to bones and connective tissue, such as the thoracolumbar fascia, the nuchal and sacrospinous ligament and the linea alba (Table 2). In addition, we also included several connections between bones which barely move relative to each other, for example the os coxae, os ilium, os ischii and ligamentum sacrospinale which together form the pelvis (*25*).

To evaluate how the anatomical muscle network is shaped by physical connections that are included, we compared the current network to the anatomical network presented in Murphy *et al.* (*19*), which is based on the Hosford Muscle table. We used the bipartite network provided in the Supporting Information (https://doi.org/10.1371/journal.pbio.2002811.s016) and selected the 36 muscles used in the current study. Compared to the bipartite network used in the current study, the bipartite network in Murphy *et al.* includes a larger number of individual bones (e.g. all 12 ribs are listed individually), while some connective tissues (fascia thoracolumbalis, ligamentum sacrospinale and linea abla) are not included. We then defined the muscle-centric network as the one-mode projection.

The resulting adjacency matrix reveals a number of differences (Fig. S1). The Kerkman network includes several connections that are absent in the Murphy network. For example, RF and AL are connected in the Kerkman network but not in the Murphy network, because RF is attached to the os ilium, whereas the AL is attached to the os pubis. These are considered as separate bones in the Murphy network, whereas they are considered as a part of the pelvis in the current paper and in Diogo *et al.* (*25*). There are two connections (PMA-BB and LD-BB) that are present in the Murphy network but not the Kerkman network. This is because the BB is considered to be connected to the humerus in the Murphy but not in the Kerkman network. The density of Murphy network is lower (0.18) than density of the Kerkman network (0.27), whereas the modularity of Murphy network is higher (Q = 0.46) than the Kerkman network (Q = 0.39).

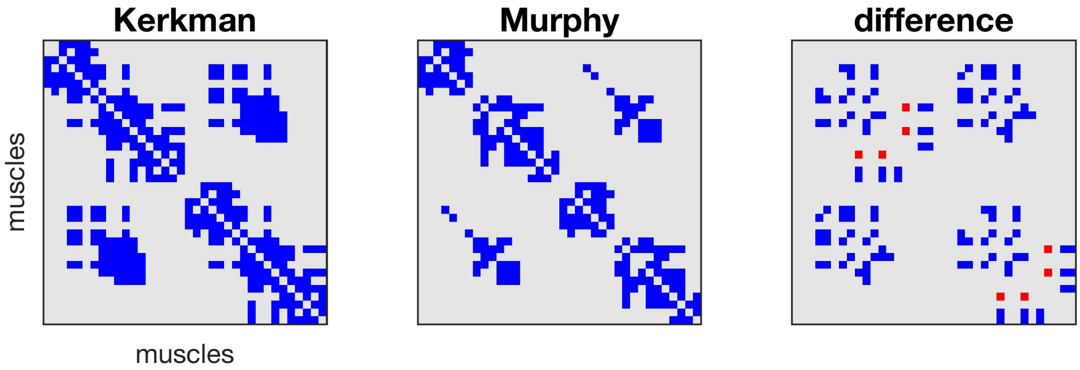


**Figure S1. The adjacency matrix of anatomical muscle networks**. The left panel shows the adjacency matrix of the muscle-centric network used in the present study. The middle panel shows the adjacency matrix used in Murphy *et al.* (*19*). The right panel shows the difference in the adjacency matrices. Blue squares indicate additional connections between muscles in the Kerkman network and red additional connections in the Murphy network.

**Spinal nerve network**

Muscles are innervated by axonal terminals of motor neurons located in the ventral column of the spinal cord and afferent nerve fibres transmit sensory information from muscle spindles and Golgi tendons back to the dorsal column of the spinal cord. These efferent and afferent connections are part of the spinal nerve that connects the spinal cord and the muscle. In the human body there are 31 pairs of spinal nerves, one on each side of the vertebral column. Different spinal nerves innervate different muscles (e.g., arm muscles are innervated by cervical nerves, while leg muscles are innervated by lumbar nerves) and a single spinal nerve innervates a number of different muscles – a myotome (e.g., D and BB are both innervated by cervical spinal nerve 5). Shared innervation by spinal nerves may be a potential factor that contributes to the common input different muscles receive, although spinal fMRI may provide a more detailed map of the functional connectivity in the spinal cord (*47*).

We constructed a bipartite network between muscles and spinal nerves based on gross human anatomy (Table S1). A muscle-centric network was obtained using the one-mode projection. The resulting adjacency matrix reveals a number of three communities, two for the upper body and one for the legs (Fig. S2). The upper- and lower-body modules are unconnected as there is no overlap in innervation between muscles in the upper and lower body: All upper body muscles are innervated by nerves L1 and above while all lower body muscles are innervated by nerves below L1. The spinal nerve network has a density of 0.36 and a modularity of 0.46.

**Table S1. Spinal nerve innervation**

| Muscle | Spinal nerves |
|---|---|
| TA | L4-S1 |
| GM | S1-S2 |
| SOL | S1-S2 |
| RF | L2-L4 |
| BF | L5-S3 |
| VL | L2-L4 |
| AL | L3-L4 |
| EO | T1-L1 |
| PMA | C5-T1 |
| SMA | C2-C3 |
| LO | C1-T12 |
| LD | C6-C8 |
| TZ | C3-C4 |
| D | C5-C6 |
| BB | C5-C6 |
| TRB | C5-C6 |
| ED | C6-C8 |
| FDS | C7-T1 |

Innervations are based on gross human anatomy as described in Martini *et al.* (*2*).

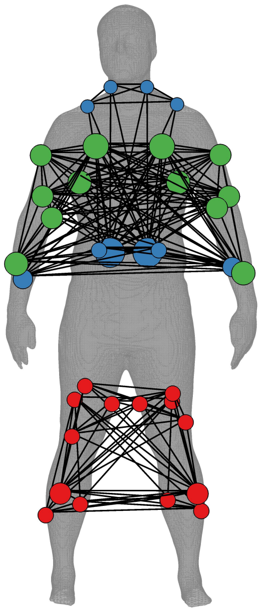

**Figure S2. Community structure of the spinal nerve network.** Spatial representation of spinal nerve muscle network displayed on the human body (*26*). The size of each node represents the number of other nodes it is connected to. Colours indicate different communities.

### Weighted functional networks

Intermuscular coherence was used to estimate the weights of the functional muscle networks. We then thresholded the weights to obtain a minimally connected binary network and facilitate the comparison to the binary anatomical network. The Louvain algorithm can also be used to determine the community structure of weighted networks.

Performing the analysis directly on the weighted networks may provide additional information about functional coordination. We therefore assessed the community structure of the weighted functional network for each frequency component separately. The weights were averaged across the nine conditions and the community structure was assessed using consensus clustering (Fig. S3). We observed three, three, five, and six modules and a modularity of 0.28, 0.27, 0.34, and 0.28 for frequency components 1 to 4, respectively. The results showed increased modularity at higher frequencies and are qualitatively similar to the results on binary networks. Future studies may extend the multiplex modularity analysis (*24*) to weighted networks to enable quantitative comparisons between binary and weighted multiplex networks.

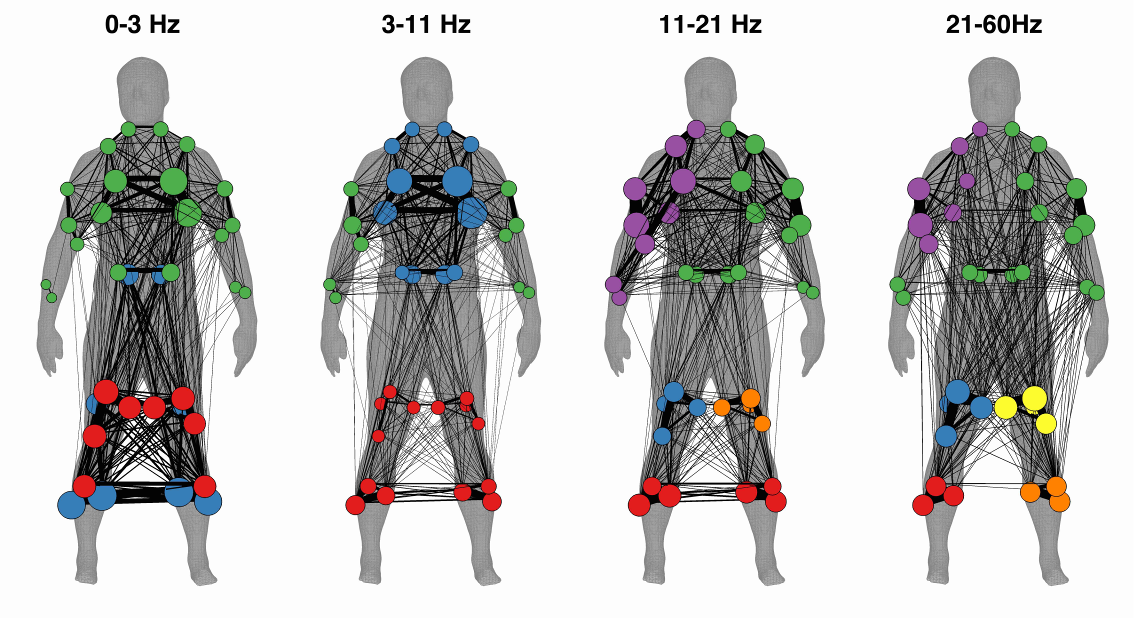


**Figure S3. Community structure of weighted functional network.** The community structure was determined separately for each frequency component (0-3, 3-11, 11-21, and 21-60 Hz). The size of each node represents the number of other nodes it is connected to and the width of the edges the average weights. Colours indicate different communities.
